# The RNA-binding protein Adad1 is necessary for germ cell maintenance and meiosis in zebrafish

**DOI:** 10.1101/2022.12.21.521539

**Authors:** Kazi Nazrul Islam, Anuoluwapo Ajao, Katrin Henke, Kellee R. Siegfried

## Abstract

The double stranded RNA binding protein Adad1 (adenosine deaminase domain containing 1) is a member of the <u>a</u>denosine <u>d</u>eaminase <u>a</u>cting on <u>R</u>NAs (Adar) protein family with germ cell-specific expression. In mice, Adad1 is necessary for sperm differentiation, however its function outside of mammals has not been investigated. Here, through an N-ethyl-N-nitrosourea (ENU) based forward genetic screen, we identified an *adad1* mutant zebrafish line that develop as sterile males. Further histological examination revealed complete lack of germ cells in adult mutant fish, however germ cells populated the gonad, proliferated, and entered meiosis in larval and juvenile fish. Although meiosis was initiated in *adad1* mutant testes, the spermatocytes failed to progress beyond the zygotene stage. Thus, Adad1 is essential for meiosis and germline maintenance in zebrafish. We tested if spermatogonial stem cells were affected using a label retaining cell (LRC) assay and found that the mutant testes had fewer LRCs compared to wild-type siblings, suggesting that failure to maintain the spermatogonial stem cells resulted in germ cell loss by adulthood. To identify potential molecular processes regulated by Adad1, we sequenced bulk mRNA from mutants and wild-type testes and found mis-regulation of genes involved in RNA stability and modification, pointing to a potential broader role in post-transcriptional regulation. Our findings suggest that Adad1 is an RNA regulatory protein required for fertility through regulation of spermatogonial stem cell maintenance in zebrafish.

**Author Summary:** Infertility is a serious problem for millions of couples who wish to have children. Globally more than 10% of couples suffer from infertility due to genetic, epigenetic, and environmental factors. Among these about 50% of cases occur due to genetic factors such as aneuploidy and genetic mutations affecting development of the gametes (i.e. sperm and eggs). Although many genes are known to be involved in germ cell development, genetic causes of infertility are still largely unexplained. Therefore, it is imperative to investigate genes involved in reproductive processes. In this study, we report that the *adad1* gene is essential for germ cell maintenance and fertility in zebrafish. Our analysis of zebrafish *adad1* mutants demonstrates that it is required for maintenance of the germline stem cells and for completion of meiosis. This is in contrast to mouse *Adad1*, which functions later in gamete development to regulate differentiation of haploid sperm. Our work on zebrafish *adad1* has uncovered previously unknown roles of *adad1* function in germline stem cell maintenance.

## Introduction

Reproduction in multicellular organisms generally depends on the differentiation of gametes from germline stem cells. In many organisms, fertility is maintained throughout life through continuous replenishment of the germ cells. The testis of most animals, including zebrafish and humans, contain spermatogonial stem cells (SSCs) which enable them to maintain continuous sperm production. SSCs are undifferentiated male germ cells which have the ability to self-renew in order to maintain a stem cell population, as well as to produce progenitor cells that differentiate into sperm (1,2). How the SSC population is established and maintained throughout life is not completely understood and is critical to understanding infertility.

The process of spermatogenesis is similar among vertebrates, except that where anamniote vertebrates (fishes and amphibians) have a cystic mode of spermatogenesis, amniotes (reptiles, birds, and mammals) have non-cystic type spermatogenesis (3). In the zebrafish, which have cystic spermatogenesis, each cyst is derived from a single spermatogonial cell that is associated with a Sertoli cell (4). Both cell types divide to produce a cyst of synchronously differentiating germ cells surrounded by Sertoli cells. Generally, in fishes, single spermatogonial cells (A_single_) have been assumed to be the SSCs. Although it is likely that A_single_ are SSCs, it is possible that additional spermatogonial type A cells, also have stem cell capacity, as has been shown in mice (1,5,6). In mouse testes spermatogonia type A_single_, A_paired_, and A_aligned_ can all act as SSCs – A_paired_ and A_aligned_ after fragmentation into A_single_ (7). In support of this, in zebrafish, the germ line stem cell marker, *nanos2*, is expressed in single spermatogonia and spermatogonia in cysts of up to four cells suggesting that A_single_ – A_4_ are the SSCs in zebrafish (8,9).

Zebrafish spermatogonia cells enter meiosis after nine rounds of mitotic divisions (4). Meiotic prophase I can be sub-divided into leptotene, zygotene, pachytene, and diplotene stage (10). In the leptotene stage, chromosomes start to condense and synaptonemal complex protein Sycp3 starts to associate with chromosome ends near the telomeres. During the late leptotene to early zygotene stage, all telomeres are associated with the nuclear envelope and cluster to one side of the nuclear envelope to form a zyotene bouquet. It has been hypothesized that meiotic bouquet formation facilitates pairing of homologous chromosomes. In zygonema, Sycp1 begins to associate with chromosomes near the telomeres, following the extension of Sycp3 along chromosomes as the zygotene stage progresses. In the pachytene stage, all homologs synapse together and begin to exchange their DNA via recombination. This stage is characterized by end-to-end association of Sycp1 and Sycp3 on the chromosomes, marking a fully formed synaptonemal complex. Homologs start separating from each other in the diplotene stage followed by the homologue segregation and cell division. Completion of meiosis II and spermiogenesis gives rise to haploid sperm. Disruption of meiosis or spermiogenic phases can cause failed or abnormal gamete formation and infertility.

Post-transcriptional RNA regulation is critical for many aspects of spermatogenesis. RNA binding proteins (RBPs) form ribonucleoprotein complexes which are important for post-transcriptional regulation of RNAs such as translational control, RNA stability, RNA biogenesis, and modifications of RNAs (11). During spermatogenesis, RBPs play critical roles in all stages of spermatogenesis. For example, RBPs function to protect genome integrity through destabilization of transposon RNAs, for translational regulation of mRNAs encoding proteins that function during transcriptionally silent stages (i.e. during spermiogenesis), and to regulate RNA stability during transition from one spermatogenic stage to the next (12). The adult human testis has the highest number of tissue-specific RBPs suggesting a particularly important role of post-transcriptional RNA regulation as a means to control processes specific to the male germ line (13). As such, understanding the function of germline-specific RBPs is critical to uncovering developmental mechanisms that govern spermatogenesis.

Here, through an ENU mutagenesis screen we identified a zebrafish mutant line (*t30103*) that failed to maintain germ cells during adulthood. These mutants had germ cells in larval and juvenile stages but rapidly lost the germ cells at adulthood. Through positional cloning and whole exome sequencing, we identified a missense mutation in the *adenosine deaminase domain containing 1* (*adad1*) gene that was responsible for the germ cell loss in this mutant line. The *adad1* gene encodes a germ cell-specific dsRNA binding protein and is conserved in vertebrates. The Adad1 protein has a deaminase domain similar to Adar (adenosine deaminase acting on RNA) RNA editases, however key catalytic residues are missing in Adad1 and RNA editing activity has not been detected suggesting that Adad1 regulates RNAs through a different mechanism (14). We found that zebrafish *adad1* mutants had defects in SSC maintenance as well as in meiosis indicating that *adad1* functions in multiple stages of spermatogenesis. Furthermore, transcriptomics analysis showed that RNA processing and modification pathways were disrupted in the absence of Adad1 function. Our work identified Adad1 as a putative RNA regulatory protein necessary for germline maintenance and fertility in zebrafish.

## Results

### *adad1* is necessary for germline maintenance in zebrafish

To identify unknown regulators of germline development, we conducted an ENU based mutagenesis screen and identified a zebrafish mutant line (named *t30103*) which developed only as males and failed to maintain germ cells during adulthood (Fig 1). To identify candidate mutations that cause this phenotype, we performed whole exome sequencing and genetic mapping. Analysis of the exome sequencing data identified two candidate mutations that were genetically linked to the *t30103* phenotype. These two mutations were located on chromosome 14 of the zebrafish genome, approximately 1 Mbp apart from each other. One mutation was found in the *slc34a1a* (*solute carrier family 34 member 1*) gene, which would result in a change of amino acid 423 from Isoleucine to Asparagine (*slc34a1a^I423N^*). The other mutation affected the *adad1* (*adenosine deaminase domain containing 1*) gene and resulted in amino acid 392 to change from Methionine to Lysine (*adad1^M392K^*) (Fig 1A and S1 Fig). Expression of both *slc34a1a* and *adad1* was detected in zebrafish testes by RT-PCR, therefore both candidate mutations could potentially result in testes defects (Fig 2 and S2 Fig).

**Fig 1.**
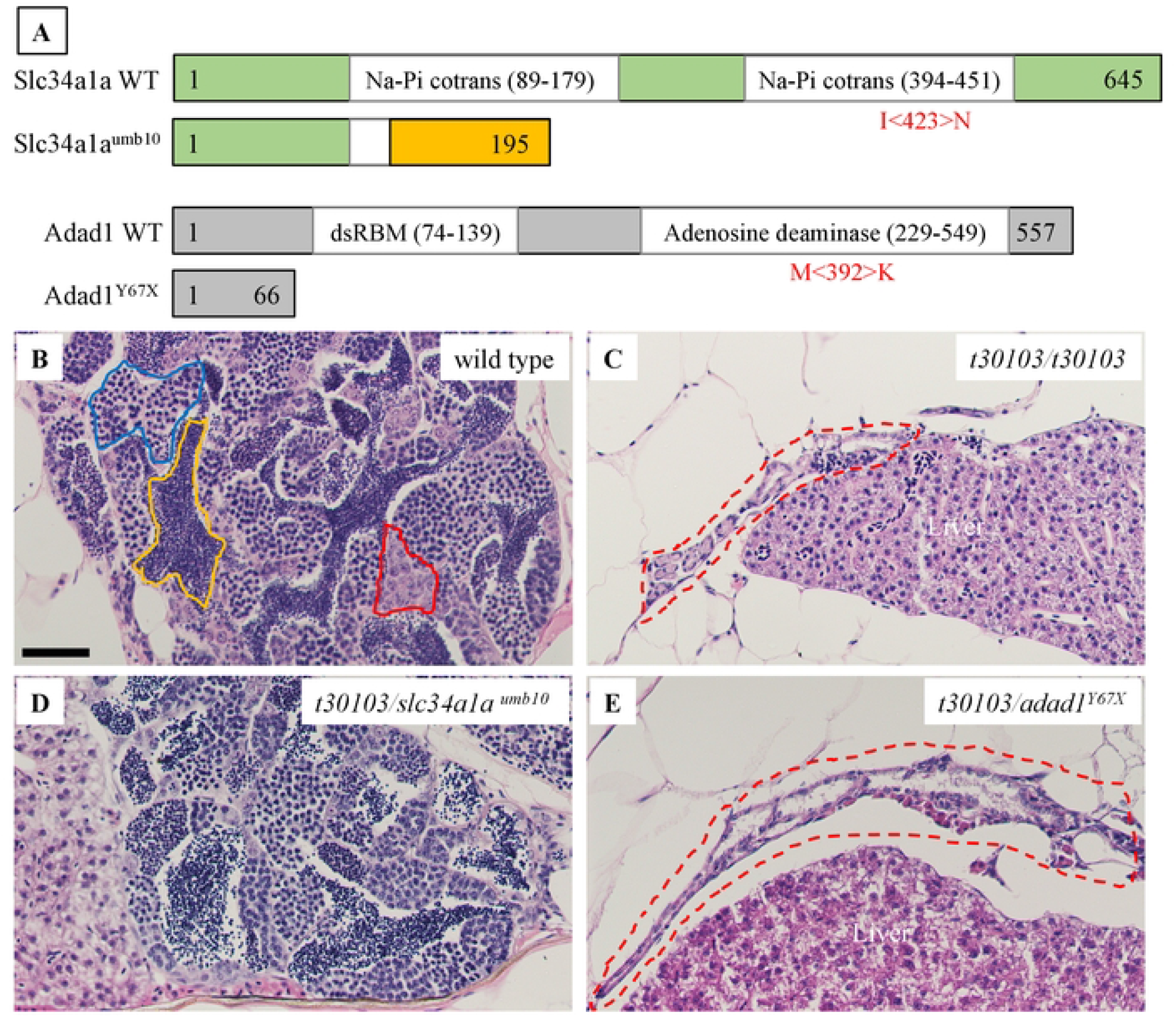
Mutations disrupting the *adad1* gene cause germ cell loss in adult zebrafish. A: Graphical representation of Slc34a1a and Adad1 wild-type and mutant proteins. Missense mutations are shown in red below wild-type protein schematics in the approximate position they occur in the protein. Representative images of truncated proteins resulting from nonsense mutations are shown. The frameshifted region of *slc34a1a^umb10^* is colored orange. Numbers denote the number of amino acids in each protein. Abbreviations: Na-Pi cotrans (sodium dependent phosphate co-transporter domain), dsRBM (double-stranded RNA binding motif), WT (wild type). B-E: Hematoxylin-eosin staining of testes from adult wild type (B), *t30103* homozygous mutant (C), *slc34a1a^umb10^* trans-heterozygous to *t30103* (D), and *adad1^Y67X^* trans-heterozygous to *t30103* (E). In the wild-type testis spermatogonia, spermatocytes, and mature sperm are marked by red, blue, and yellow outlines, respectively. The *t30103* homozygous and *adad1^Y67X^/t30103* trans-heterozygous testes are small and lack germ cells (red dotted outlines in C and E). Conversely, there is no visible difference between wild-type and *slc34a1a^umb10^/t30103* trans-heterozygous testes. Scale bar: 50 μm.

**Fig 2.**
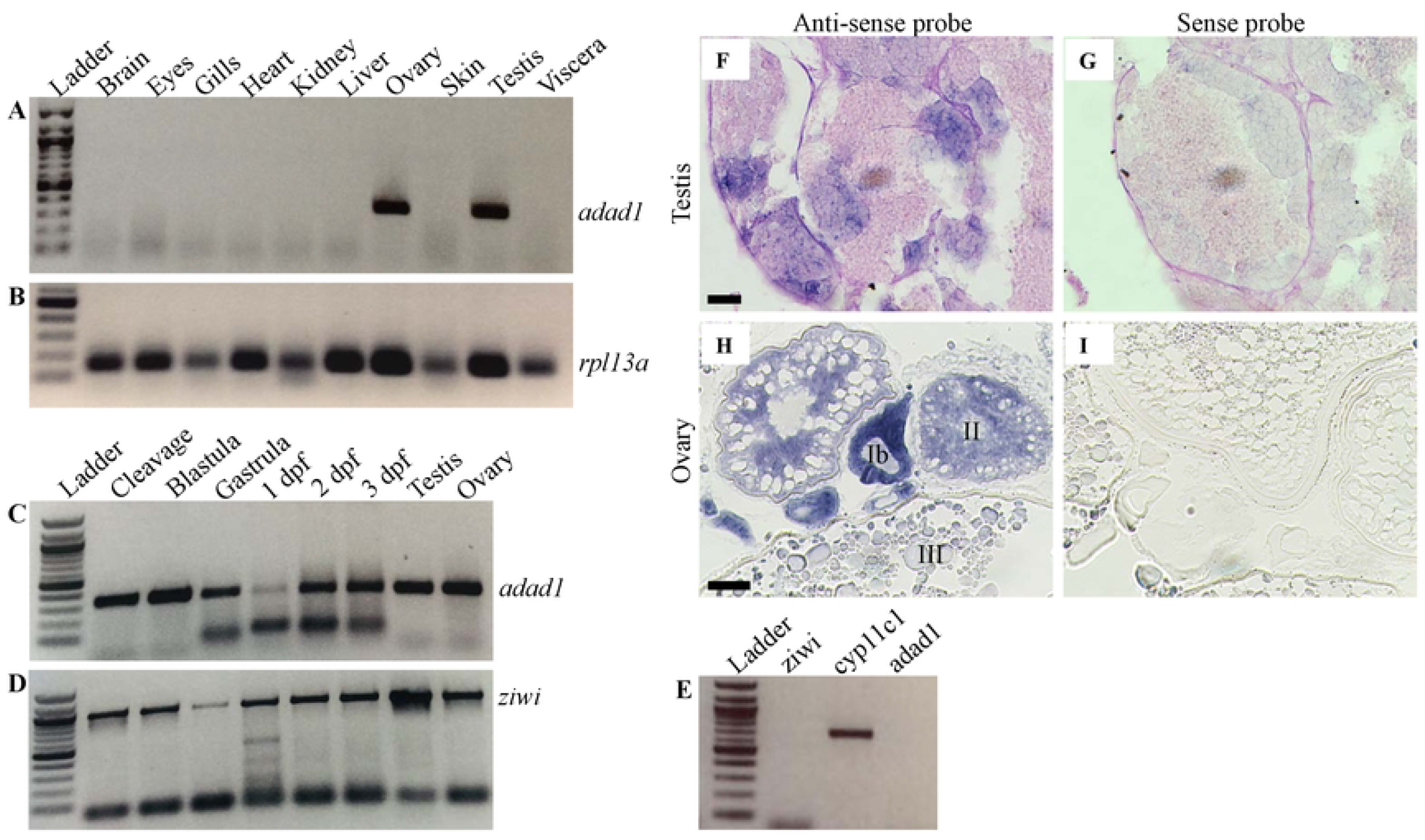
*Adad1* mRNA is expressed in both sexes and is germline specific. A-D: RT-PCR assaying *adad1* expression in wild-type tissues. RT-PCR on adult tissues demonstrates that *adad1* is expressed only in ovary and testis (A), whereas the control gene *rpl13a* is expressed in all organs tested (B). During embryonic and larval development *adad1* is expressed in all stages tested (C), similar to the germ cell marker *ziwi* (D). E: RT-PCR on testes lacking germ cells revealed that *adad1* is not expressed in somatic cells of the testis. *Ziwi* expression was not detected confirming that germ cells are not present while the Leydig cell marker gene *cyp11c1* confirmed the presence of testes tissue. F-I: *In situ* hybridization (ISH) showing *adad1* is expressed in germ cells of adult ovaries and testes. Testes were counterstained with periodic acid-Schiff. Scale bars: 20 μm for testes and 50 μm for ovaries.

To identify the causative mutation of the *t30103* line, we performed phenotypic and complementation analysis with independently derived mutations affecting each candidate gene (Fig 1A). Using CRISPR/Cas9 genome editing targeting *slc34a1a*, we generated a 17 bp deletion that caused a frameshift and premature stop codon after amino acid 195 (*slc34a1a^umb10^*), which would disrupt the transmembrane domains of the encoded channel protein (Fig 1A and S2 Fig). We obtained the *adad1^sa14397^* mutant line from the Zebrafish International Resource Center. This line carries an ENU-induced nonsense mutation at codon 67 and will be referred to as *adad1^Y67X^* henceforth (Fig 1A and S1 Fig). We performed genetic complementation analysis to test which *t30103* candidate mutation caused the germ cell defects. The *slc34a1a^umb10^* homozygous fish and *t30103/slc34a1a^umb10^* trans-heterozygotes were adult viable, developed as both males and females, and the adult males had normal testes histology (Fig 1D and S2 Fig). By contrast, *t30103/adad1^Y67X^* trans-heterozygous fish were all male and displayed a tiny testis without any germ cells at adulthood, similar to *t30103* homozygotes (Fig 1C and 1E). These data confirm that the *adad1* mutation causes the testis defect in the *t30103* mutant line. We will refer to the *adad1^t30103^* allele as *adad1^M392K^* and *adad1^sa14397^* as *adad1^Y67X^* based on their effects on the encoded protein to more easily distinguish the two alleles.

### Zebrafish *adad1* is expressed in both sexes and is germline specific

To test which organs express the *adad1* gene, we performed RT-PCR using cDNA from brains, eyes, gills, hearts, kidneys, livers, ovaries, skin, testes, and viscera. The RT-PCR results showed that *adad1* is gonad specific, as no other organs besides testes and ovaries expressed this gene (Fig 2A-B). We also asked whether *adad1* is germ cell specific in testes by performing RT-PCR on cDNA generated from testes devoid of germ cells. *Adad1* was not detected in germ cell depleted testes whereas the Leydig cell-expressed gene *cyp11c1* was detected (Fig 2E). This result suggests that *adad1* is germ cell specific in male zebrafish. We could not perform a similar experiment with germ cell-depleted ovaries because zebrafish can only develop as males when lacking germ cells (15,16). To ask whether *adad1* is expressed during embryonic or early larval stages, we performed RT-PCR using cDNA from whole embryos and larvae of the cleavage, blastula, gastrula, 1 day post fertilization (dpf), 2 dpf, and 3 dpf. *Adad1* expression was detected in all stages tested (Fig 2C and 2D). Next, to see which type of germ cells expressed *adad1* mRNA, we performed *in situ* hybridization (ISH) on testis and ovary sections of adult zebrafish. In testes, *adad1* was expressed in both spermatogonia and spermatocytes, but not in the mature sperm (Fig 2F and 2G). The ovary ISH showed high *adad1* expression in stage Ib and stage II oocytes, and faint expression in later stage oocytes, with no apparent expression in somatic cells (Fig 2H and 2I). These data show that zebrafish *adad1* is expressed in germ cells of both sexes.

To enhance our understanding of Adad1, we made an antibody against zebrafish Adad1 protein. We tested specificity of the antibody through western blot and immunofluorescence (IF) using mutant and wild-type testes (S3 Fig). In both experiments, mutant testes were collected from juvenile fish prior to the time at which germ cells are absent, as evident from detection of the germ cell-specific protein Vasa (S3 Fig). By western blot we detected a predicted 62 kDa sized protein in wild-type testes extracts, whereas no bands were detected in testes extracts from the *adad1^Y67X^* mutants (S3E Fig). Similarly, no protein was detected by IF on *adad1^Y67X^* mutant testis while protein was visible in wild-type testes (Fig 3 and S3A-D Fig), indicating that the antibody is specific for Adad1. Analysis of IF performed on wild-type adult testis and ovary sections showed germ cell-specific expression of Adad1 protein in both sexes (Fig 3). The IF on ovary sections demonstrated that Adad1 is highly expressed in stage Ib and stage II, and has little to no expression in later stage oocytes (Fig 3A-D). In the testis, Adad1 was expressed highly in spermatogonia and moderately in the spermatocytes but undetectable in the sperm (Fig 3E-H). Interestingly, we observed both nuclear and cytoplasmic localization of Adad1 in early spermatogonia (1 cell and cyst of up to 4 cells), whereas later stage spermatogonia and spermatocytes showed only cytoplasmic localization (Fig 3I-L). This observation suggests that Adad1 may play a unique role in early stage spermatogonia, which includes the putative spermatogonial stem cells, compared to later stage germ cells.

**Fig 3.**
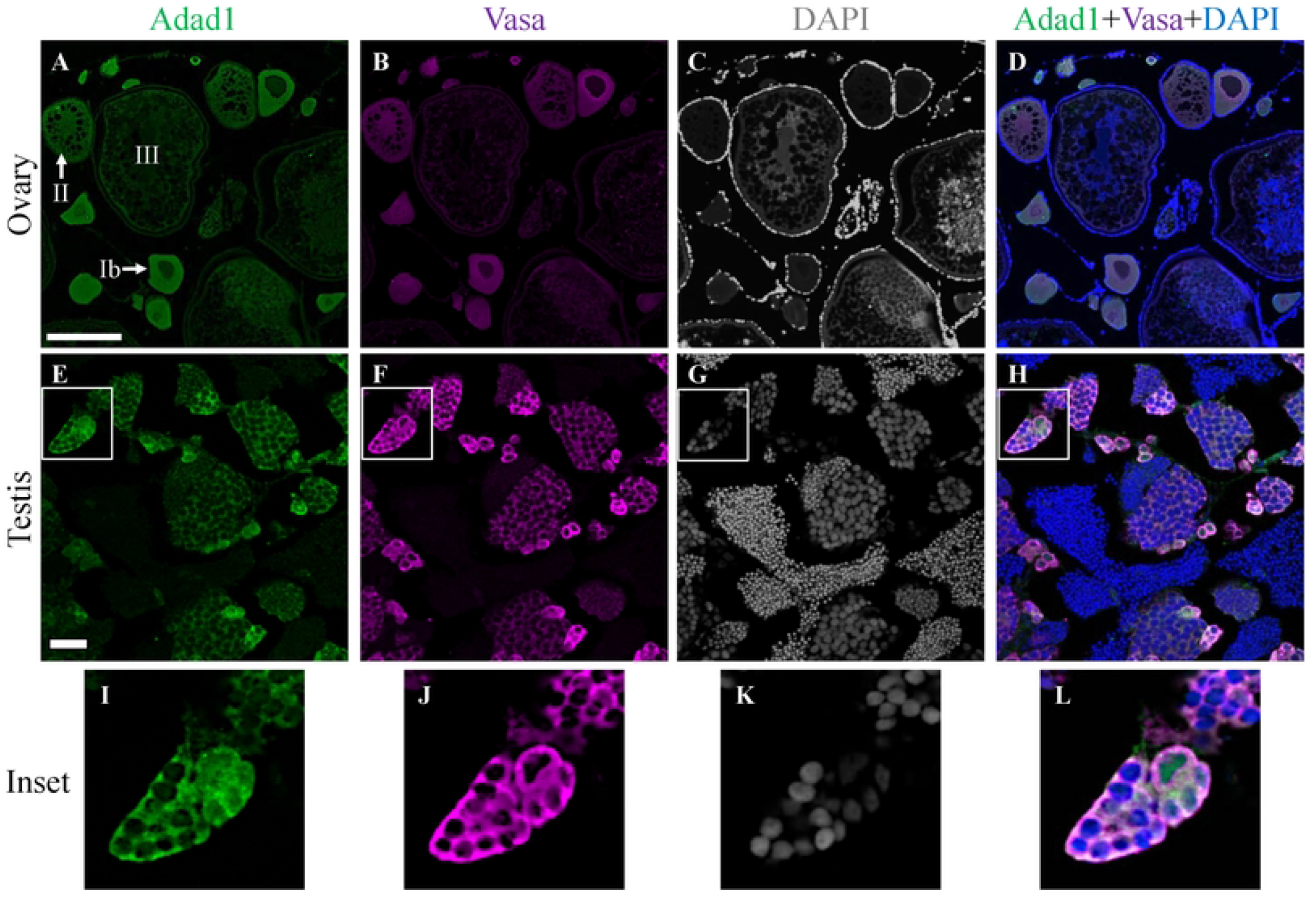
Adad1 protein expression in adult ovaries and testes. A-D: In the ovary, Adad1 is expressed in the cytoplasm of stage Ib and stage II oocytes (A) similar to the Vasa protein (B). E-L: In the testis, Adad1is expressed in the cytoplasm of spermatogonia and spermatocytes but not in mature sperm (E), overlapping with Vasa expression (F). Magnification of the insets shows that Adad1 is both nuclear and cytoplasmic in single spermatogonia and spermatogonia in cysts of 2-4 cells. Scale bars: 100 μm for ovary and 20 μm for testis.

### Adad1 is essential for spermatogenesis and oogenesis in zebrafish

To identify the developmental stage at which *adad1* mutants lost their germ cells, we did histology on gonads from juvenile (45 and 60 dpf) and adult (90 dpf) mutant and wild-type fish (Fig 4). Histological analysis showed that *adad1* mutants had germ cells at 45 dpf similar to the wild types (Fig 4A and 4B). At 60 dpf, both mutant and wild-type testes had abundant germ cells, however wild-type fish had some mature sperm, but mutant testes lacked sperm (Fig 4C and 4D). To rule out the possibility of developmental differences in *adad1* mutants due to growth delays, we measured the standard length of the fish prior to fixation and found them to be comparable between mutants and wild types. These results suggest that *adad1* is dispensable for embryonic and early larval germ cell development and for the initiation of spermatogenesis, but is required to produce mature sperm. The histology of testes from 90 dpf mutant fish were completely devoid of germ cells in contrast to wild-type testes which had a robust germ cell population undergoing spermatogenesis (Fig 4E and 4F). These results indicate that *adad1* mutants lost their germ cells during the juvenile to adult stage transition.

**Fig 4.**
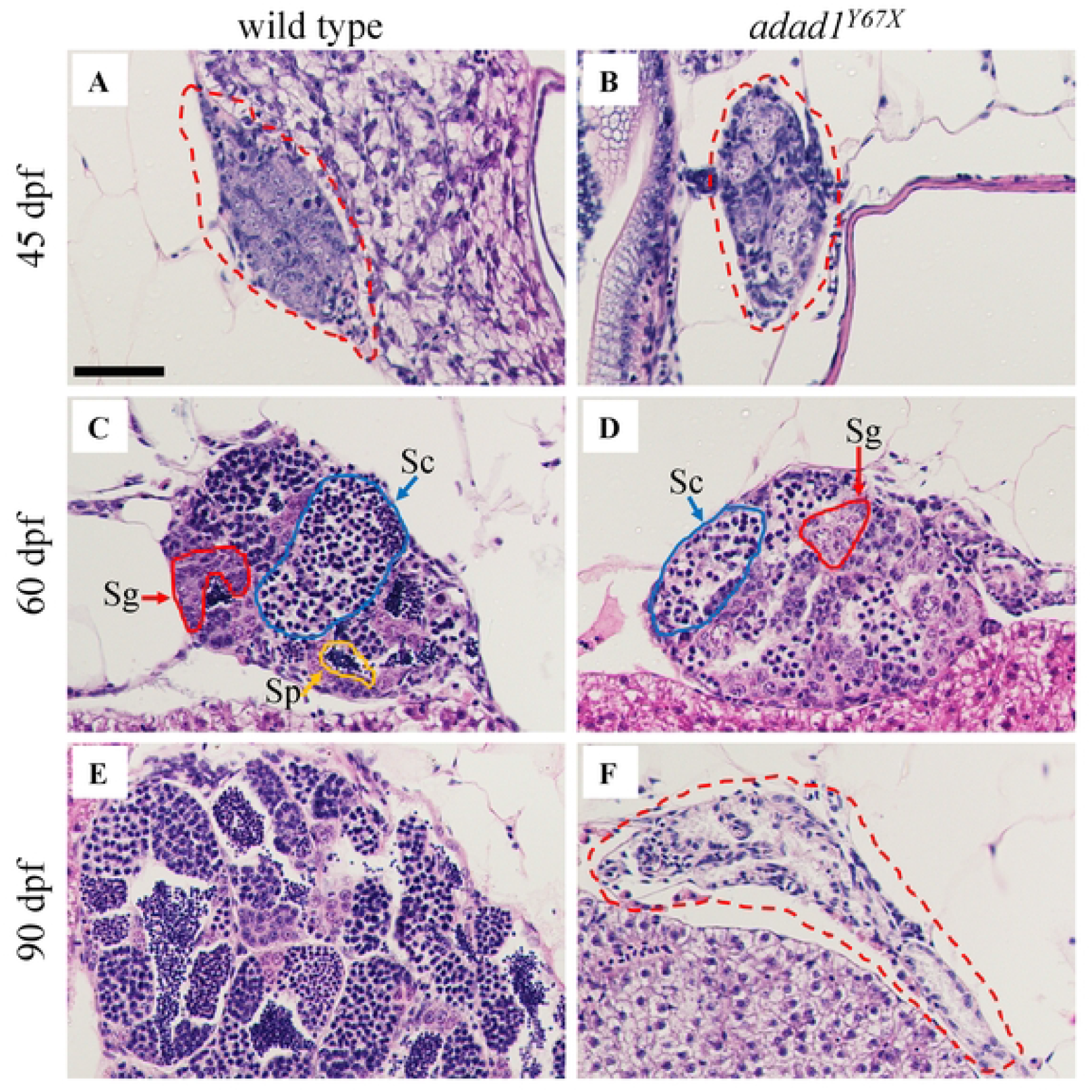
Histology of wild-type and *adad1^Y67X^* mutant gonads at 45, 60, and 90 dpf. A-B: At 45 dpf, germ cells are present in both the wild-type (A) and mutant (B) testes. C-D: At 60 dpf, spermatogonia (Sg, red outline) and spermatocytes (Sc, blue outline) are present in both wild-type (C) and mutant (D), but the mutant lacks mature sperm (Sp, yellow outline). E-F: At 90 dpf, the wild-type testis is larger with abundant sperm (E) whereas the mutant testis is small and devoid of germ cells (F). Dotted red lines outline the gonads in A, B and F. Scale bar: 50 μm.

We also tested whether *adad1* mutant males were capable of inducing spawning in females. Breeding experiments showed that *adad1* mutant males (N=3) successfully induced wild-type females to release eggs even though mutant males were unable to fertilize them (Table 1). These data suggest that *adad1* mutants have normal male mating behavior.

**Table 1.**
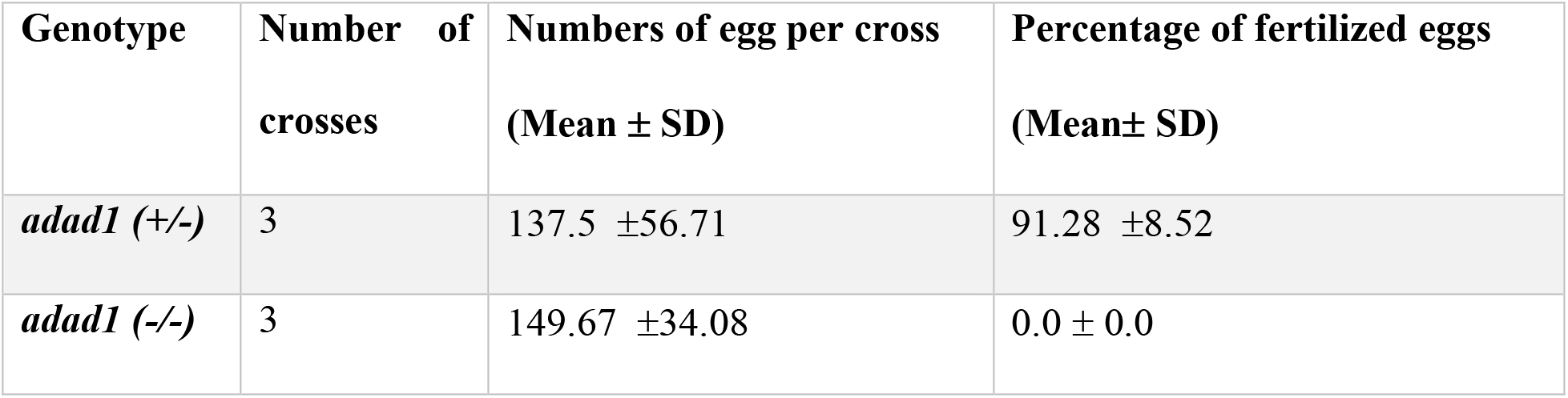
Fertility test of *adad1^Y67X^* mutant fish.

| Genotype | Number of crosses | Numbers of egg per cross<br>(Mean ± SD) | Percentage of fertilized eggs<br>(Mean ± SD) |
| --- | --- | --- | --- |
| *adad1* (+/-) | 3 | 137.5 ± 56.71 | 91.28 ± 8.52 |
| *adad1* (-/-) | 3 | 149.67 ± 34.08 | 0.0 ± 0.0 |

Since all *adad1* mutants developed as males, we wondered whether or not mutants could initiate female development. Zebrafish manifest their gonadal sex at around 30 dpf, although this can vary within different zebrafish facilities. Before this age, the zebrafish gonad remains undifferentiated and passes through a juvenile ovary-stage, where immature oocytes are present, before finally differentiating into either an ovary or testes (17). Thus, all zebrafish display a juvenile ovary prior to gonadal sex differentiation. To ask if *adad1* mutant fish could initiate oogenesis, we did histology at 28, 35, and 42 dpf. Because the precise timing of zebrafish sex differentiation can be variable, we compared histology of mutant and wild-type siblings at each time point, using wild-type siblings to determine the developmental stage of the population. Histology at 28 dpf showed that the gonads were still in an undifferentiated state and had not initiated oogenesis in either the mutant (N=6) or wild-type (N=6) fish (Fig 5A and 5B). However, histology at 35 dpf showed that all *adad1* mutants (N=6) were developing as males, whereas their wild-type siblings were developing both as males or females (Fig 5C-E). We found similar results for the 42 dpf fish (Fig 5F-H). These results suggest that *adad1* mutants failed to initiate oogenesis, therefore this gene is necessary for both female and male germ cell development.

**Fig 5.**
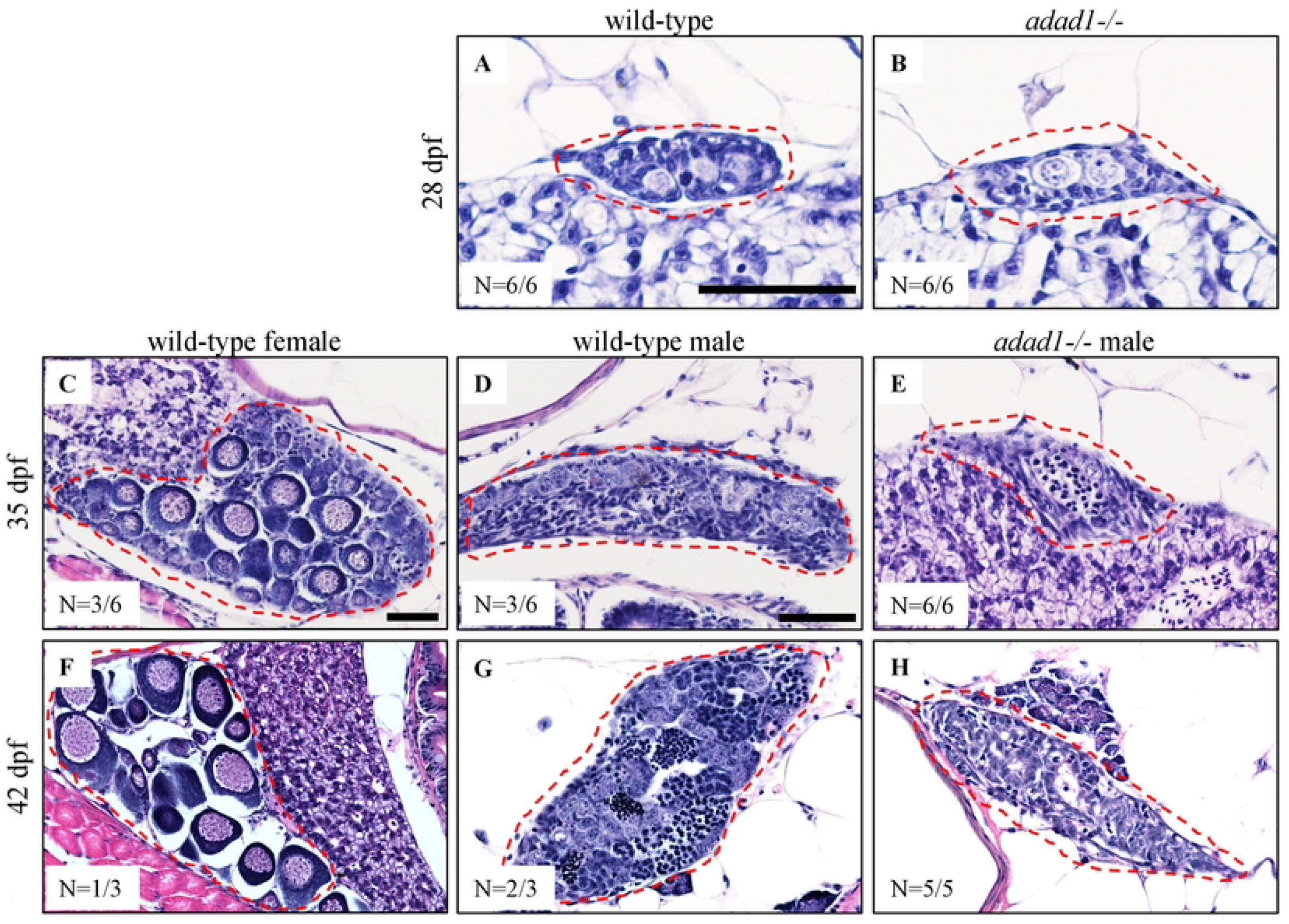
Hematoxylin-eosin staining on 28, 35, and 42 dpf wild-type and mutant gonads. A-B: At 28 dpf, gonads are undifferentiated in wild-type (A) and *adad1^Y67X^* mutant (B) fish. C-H: At 35 and 42 dpf, wild-type fish were developing as either female (C, F) or male (D, G), whereas mutant fish were developing only as males (E, H). Gonads are marked by dotted red outlines in all images. Scale bar: 50 μm.

### Adad1 is needed to complete the zygotene stage of meiosis prophase I

Since we observed that *adad1* mutants produced spermatocytes but failed to make mature sperm (Fig 4D), we asked what stage of meiosis the spermatocytes could reach. Therefore, we performed IF on testes sections and on chromosome spreads to visualize key events in meiotic prophase I. IF labeling the Synaptonemal complex protein 3 (Sycp3) on testes section revealed the presence of three distinct cell populations in both the mutant and wild-type testes (Fig 6). In wild-type testes, leptotene stage nuclei had few Sycp3 dots, early zygotene/bouquet stage nuclei had bright Sycp3 concentrated on one side, and late zygotene/pachytene stage cells had Sycp3 evenly dispersed across the nucleus (Fig 6A-C). The presence of similar cell populations in the mutants suggests that *adad1* mutant spermatocytes can reach at least the zygotene stage (Fig 6D-F). However, co-labelling with nuclear stain DAPI showed that the mutant testis lacked mature sperm, as observed in histological sections (Fig 4D and Fig 6E). To better define the meiotic stage at which *adad1* mutant spermatocytes stop developing, we did IF for Sycp1 and Sycp3 together with telomere fluorescence *in situ* hybridization (Telo-FISH) on nuclear spreads (Fig 7). In wild types, leptotene stage cells are characterized by Sycp3 expression near chromosome ends with Sycp1 either few or no visible puncta (Fig 7A-E). In the early zygotene stage cells, telomeres cluster together to form the meiotic bouquet where both Sycp1 and Sycp3 begin to load on chromosomes as homologous chromosomes start pairing (Fig 7F-J). Late zygotene stage cells have elongated Sycp1 and Sycp3 lines as synapsis extends toward the interstitial region of the chromosomes (Fig 7K-O). The pachytene stage cells have Sycp1 and Sycp3 labelling from end to end of homologous chromosomes (Fig 7P-T) (10). In *adad1* mutants, we could see leptotene stage cells similar to the wild types (Fig 7A’-E’). We also found that some *adad1* mutant spermatocytes had Sycp1 and Sycp3 loaded onto chromosomes, indicating that synapsis and pairing were initiated, characteristic of zygotene stage cells (Fig 7F’-J’). However, we did not find any mutant spermatocytes in the pachytene stage. These observations suggest that *adad1* mutant spermatocytes initiated meiotic prophase I but failed to reach the pachytene stage, therefore Adad1 is necessary to complete meiotic prophase I in zebrafish.

**Fig 6.**
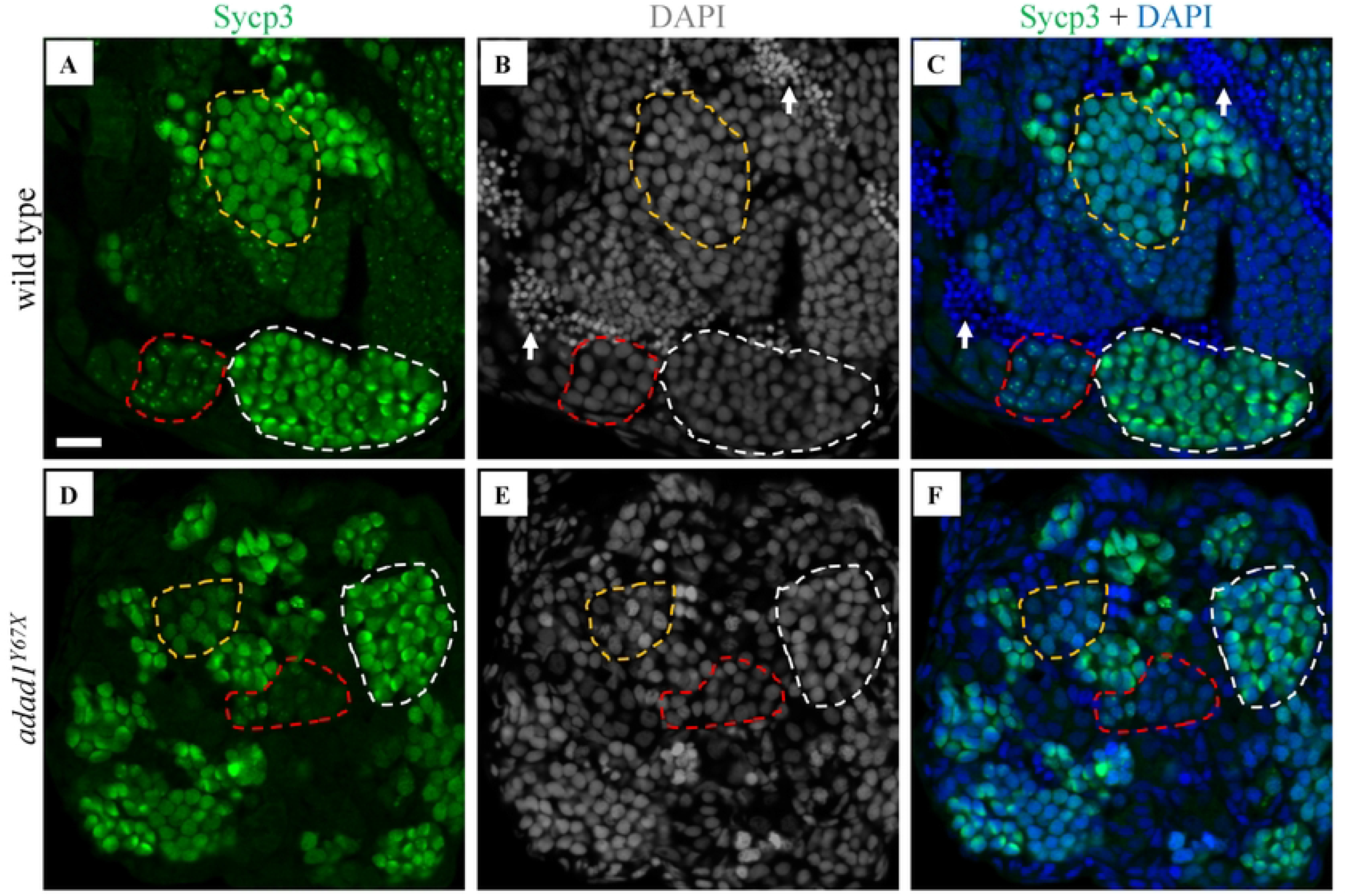
Immunostaining with meiotic marker Sycp3. A-F: Sycp3 immunofluorescence (green) and DAPI nuclear stain (white in B, E, K and blue in C, H, L) on 60 dpf wild-type (A-C) and *adad^Y68X^* testis (D-F) showed the presence of leptotene (red dotted outline), early zygotene or bouquet (white dotted outline), and late zygotene/pachytene (yellow dotted outline) stage cells in both wild-type (A-C) and *adad1* mutant (D-F) testes. Arrows point to sperm. Scale bar: 20 μm.

**Fig 7.**
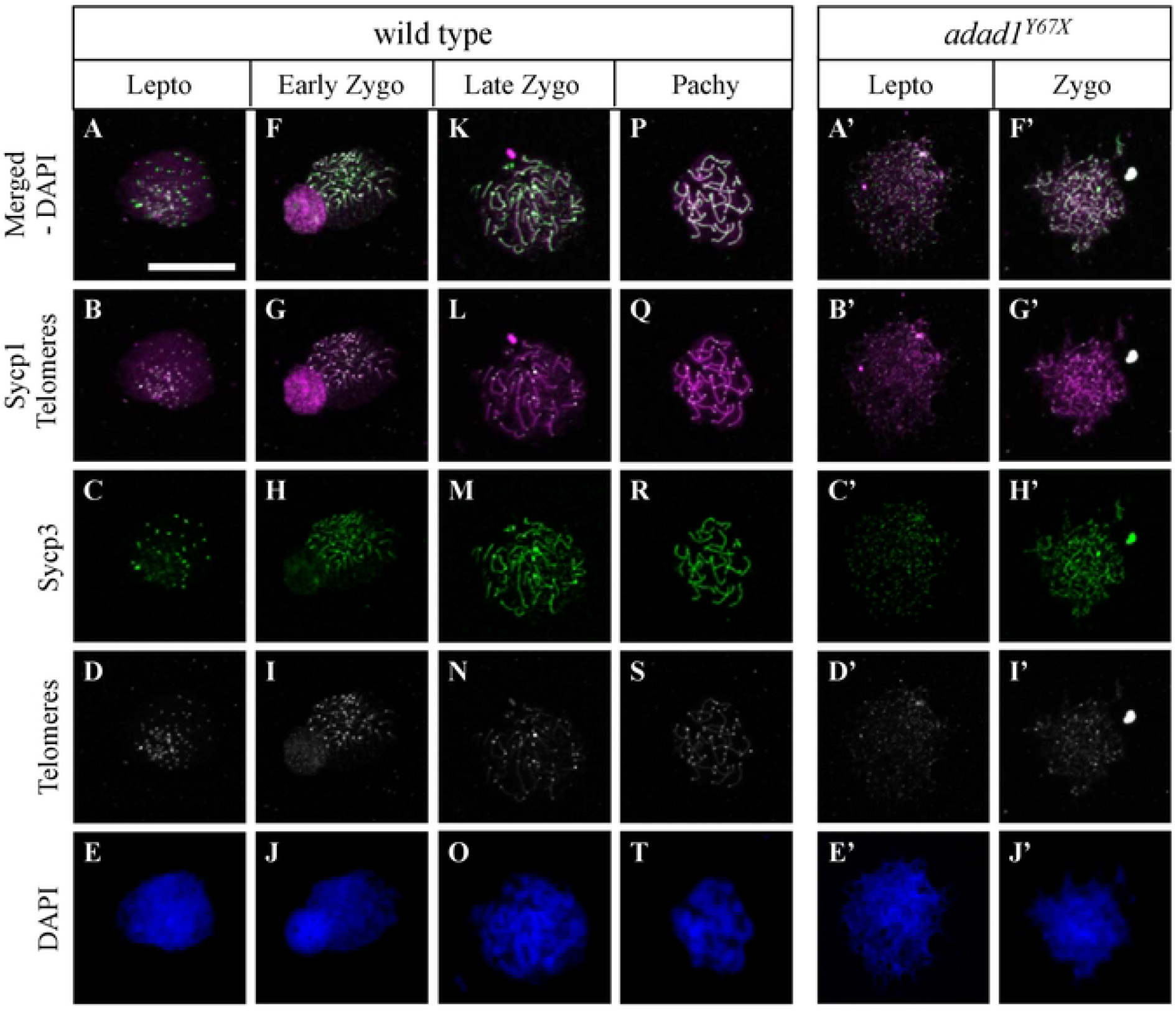
Telomere and synaptonemal complex protein labelling on meiotic chromosome spreads. A-T: Wild-type chromosome spreads have leptotene (A-E), zygotene (F-O), and pachytene (P-T) stage cells. A’-J’: Chromosome spreads from *adad1^Y67X^* testes lack pachytene stage cells. Telo-FISH (white), IF for Sycp1 (magenta) and Sycp3 (green), and DAPI nuclear staining (blue). Scale bar: 10 μm.

### Adad1 is required for spermatogonial stem cell maintenance in zebrafish

To identify the cellular processes leading to germ cell loss at the juvenile to adult transition in *adad1* mutants, we assayed cell death, cell proliferation, and stem cells. Using a TUNEL assay and cleaved-Caspase-3 staining, we found no significant difference in the number of apoptotic cells between the mutant and wild-type testes (N=6 testes per genotype per experiment) (Fig 8A-F). Interestingly, almost all cells which were positive for TUNEL and Caspase-3 were negative for Vasa, indicating that germ cells lose Vasa expression during apoptosis. To test whether cell proliferation was reduced in *adad1* mutant spermatogonia, we labelled mitotic cells using BrdU together with Vasa and Sycp3 antibody labeling (Fig 8G-P). The cells which were positive for Vasa and BrdU, but negative for Sycp3 were considered proliferating spermatogonia (mitotic germ cells in the testis). We found no significant difference in the percentage of proliferating spermatogonia between mutants and wild types (N=6) (Fig 8P). These results suggest that *adad1* is neither required to prevent apoptosis nor to regulate spermatogonia proliferation in zebrafish.

**Fig 8.**
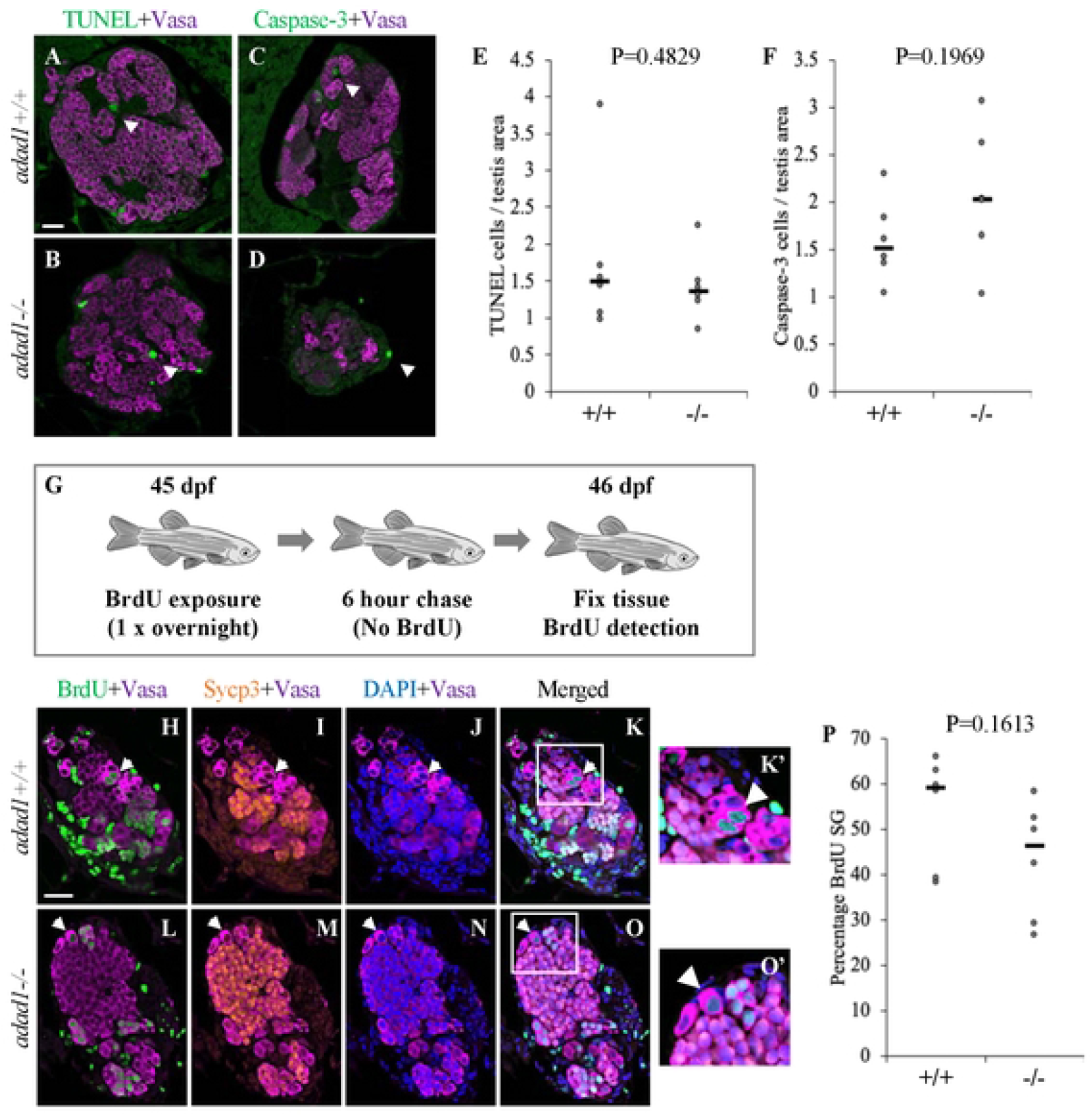
Apoptosis and proliferation are not affected in *adad1* mutants. A-F: Cell death assays in 60 dpf *adad1^Y67X^* wild-type and mutant testes. Neither TUNEL assays (A,B,E) nor cleaved-Caspase 3 immunofluorescence (B,D,F) showed a significant difference between wild-type and mutant testes. Arrowheads indicate examples of TUNEL and cleaved-Caspase3 positive cells. G-P: Cell proliferation assay in 45 dpf testes using BrdU labeling. Fish were treated overnight with BrdU then waited for 6 hours without BrdU prior to fixation (G). Immunostaining was done with BrdU, Vasa, and Sycp3 antibodies to label proliferating spermatogonia. Spermatogonia were identified as Vasa positive and Sycp3 negative (H-O). Quantification of the percentage of BrdU positive spermatogonia showed no significant difference between wild-types and mutants (P). SG = spermatogonia. Scale bar: 20 μm

We next asked whether establishment or maintenance of SSCs was defective in the *adad1* mutants by performing label retaining cell (LRC) assays, which assays for long term BrdU labeling. LRC assays have been used as a reliable method to identify quiescent stem cells in zebrafish testes (18). To identify LRCs in the testis, 32 dpf fish were treated with BrdU for three consecutive nights followed by a 25 day chase period (Fig 9A). During the chase period, all differentiating spermatogonia completed spermatogenesis and were no longer present in the testes at the end of the chase period leaving only quiescent SSCs retaining BrdU. BrdU detection was combined with Vasa, and Sycp3 antibody labeling to identify spermatogonial cells (Vasa positive, Sycp3 negative). In the wild type, LRCs were present in all testes (N=8 fish), indicating the presence of the SSCs in juvenile testes at 60 dpf (Fig 9B and 9C). On the other side, mutant testes had fewer LRCs than wild-type testes and half of the testes had no detectable LRCs (N=8 fish) (Fig 9D-F). These data show that *adad1* has a likely role in regulating SSCs.

**Fig 9.**
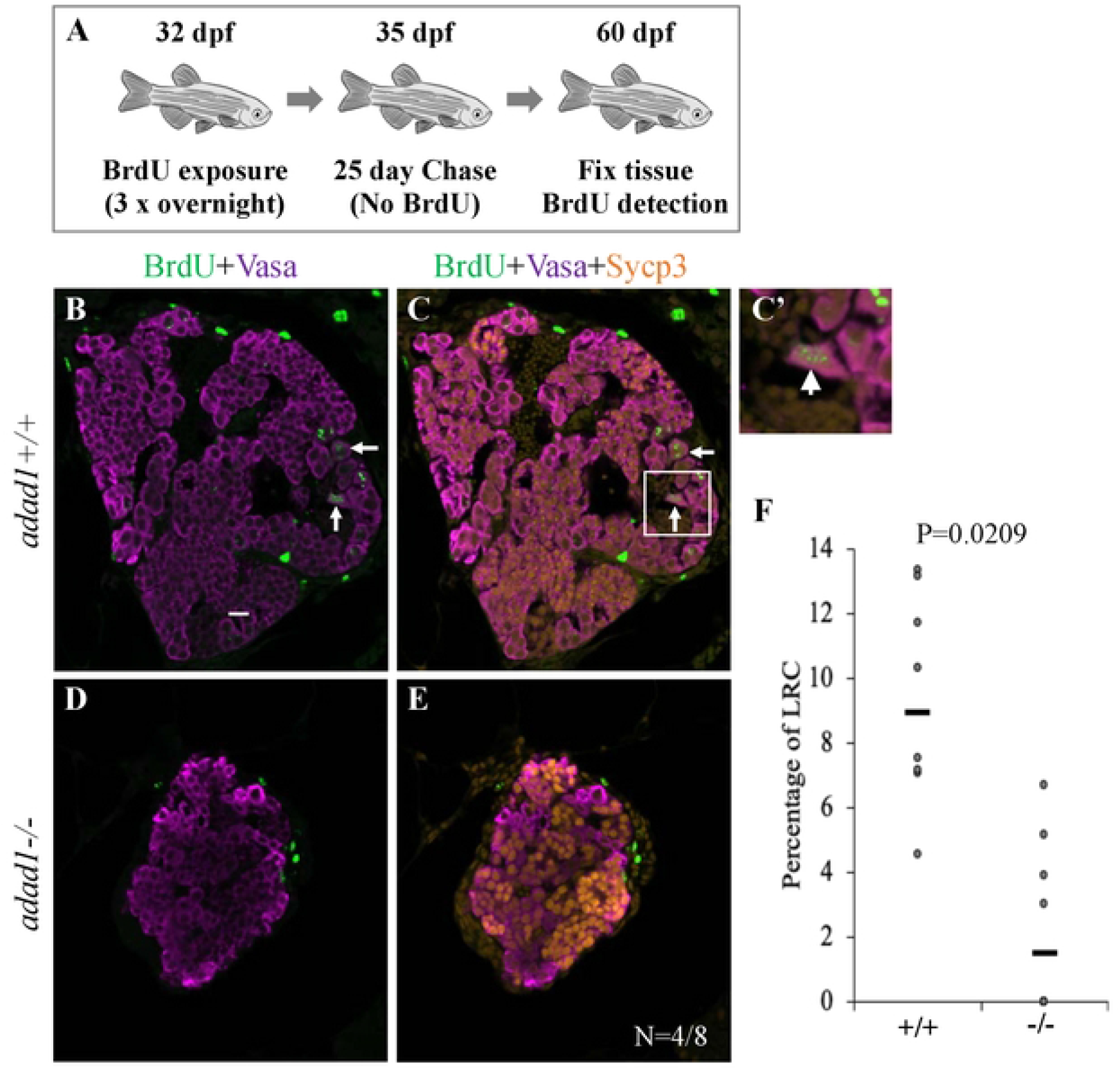
Label retaining cell assays (LRC) to identify spermatogonial stem cells. A: 32 dpf fish were treated with BrdU three times for about 16 hours per treatment (overnight) followed by a 25 day chase where fish were in water with no BrdU. At the end of the chase period, fish were fixed and processed for BrdU detection. B-F: Wild-type testes all had LRCs (B-C) with median percentage of 9.34% out of the total spermatogonia (N=8 fish) (F). Four out of eight mutant testes had no LRCs (D-E) and the median percentage of LRCs of all mutant samples was less than the wild-types (F). Arrows point to examples of LRCs. Scale bar: 20 μm.

### Adad1 is involved in regulation of DNA damage response and repair, RNA regulatory processes, and expression of specific stem cell and pluripotency genes

To identify differentially expressed genes (DEG) and potential molecular pathways through which Adad1 regulates germ cell development, we performed bulk RNA-seq from 45 dpf mutant and wild-type testes. We chose 45 dpf because the number of germ cells in the mutants are typically comparable to the wild types at this age. To identify wild-type males, we utilized germ cell-expressed GFP transgenes, which can be used to sex juvenile fish (19–21). RNA-seq was done from both the missense and non-sense mutant lines to identify common mis-regulated genes. The transcriptomic analysis identified 746 down-regulated and 697 up-regulated transcripts that were shared between both *adad1* mutant testes (Fig 10A). Interestingly, we found that genes with well-known roles in germ cell development and maintenance were generally not dysregulated in both mutant alleles (e.g. *vasa/ddx4*, *dazl*, *dnd1*, and *ziwi/piwil1*) (S1 Table). This was primarily apparent in the *adad1^Y67X^* allele, whereas *adad1^M392K^* had several such genes downregulated. We attribute this discrepancy to differences in the developmental progression of the phenotypes when the tissue was collected: based on our observations of germ cell GFP expression when collecting mutant testes, *adad1^M392K^* mutants had apparently fewer germ cells than *adad^Y67K^* testes. Therefore, we believe that the downregulated expression of multiple germ cell-expressed genes in the *adad1^M392K^* is likely a secondary effect of germ cell reduction. Pathway analysis on the shared dysregulated genes revealed that DNA repair, RNA processing and modification, noncoding RNA processing, tRNA metabolic processing pathways were downregulated in the absence of Adad1 (Fig 10C). Interestingly, we found downregulation of several genes with known roles in stem cells and pluripotency (Table 2 and S1 Table), DNA repair and meiotic recombination (Table 3 and S1 Table), meiosis (Table 4 and S1 Table), and RNA modifications (Table 5 and S1 Table) in the mutant testes. We identified *cart3*, *tdgf1*, *adad1*, *prom1b*, and *avd* as the top five downregulated genes, defined by adjusted p-value (Fig 10B). Among these *tdgf1*, which encodes a Nodal receptor, is particularly interesting because it has been shown to be involved in retaining pluripotency of mouse embryonic germ cells (22). On the other side, the top five most significantly upregulated genes are uncharacterized with the exception of *scpp7*, an unstudied apparent fish-specific calcium binding protein (Fig 10B). Overall, these results suggest that Adad1 is required for multiple cellular processes within germ cells, including DNA damage response and repair, RNA regulatory processes, and pluripotency.

**Table 2.**
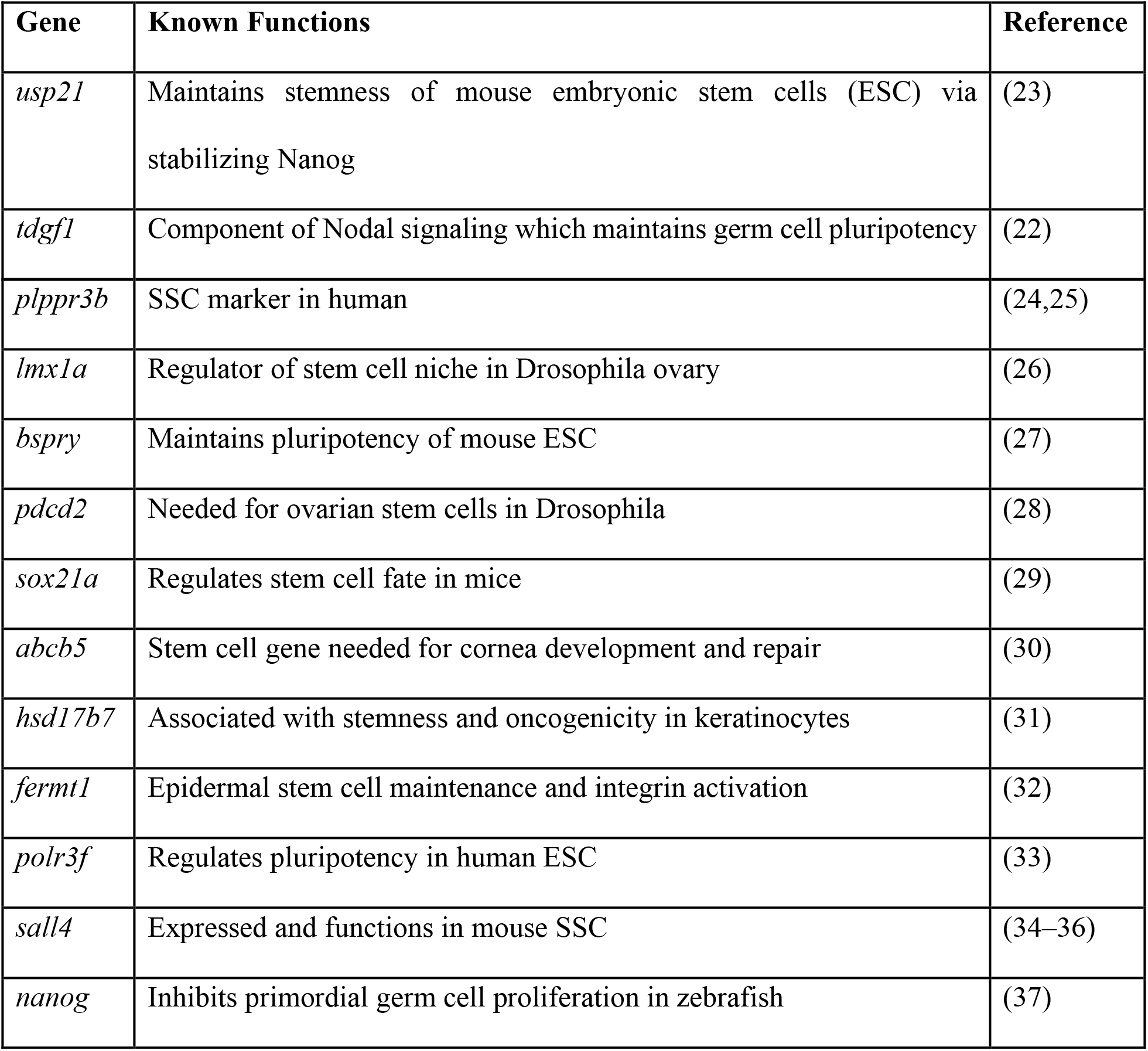
Pluripotency and stem cell genes downregulated in *adad1* mutants.

**Table 3.** DNA repair and meiotic recombination genes downregulated in *adad1* mutants.

| **Gene** | **Known Functions** | **Reference** |
| --- | --- | --- |
| *rad51ap1* | Protects cells from adverse effects of DSB | (38) |
| *hsf2bp* | Recruit recombinase to DSB sites | (39) |
| *swsap1* | Promotes early events of meiotic recombination | (40) |
| *mei4* | Regulates meiotic DSB formation | (41) |
| *helq* | Promotes Rad51 paralogue-dependent repair | (42) |
| *rad51b* | Required for DNA recombination and repair | (43) |
| *mcm8* | Needed for homologous recombination | (44) |
| *nudt1* | Oxidative DNA repair protein | (45) |
| *trip13* | Required for meiotic recombination | (46) |
| *ercc1* | Maintains integrity of germ line DNA | (47) |
| *bard1* | Localize to synaptonemal complex and regulates recombination | (48) |
| *eme1* | Holiday junction resolution and DNA repair | (49) |
| *trdmt1* | Participates in DNA damage repair | (50) |

**Table 4.**
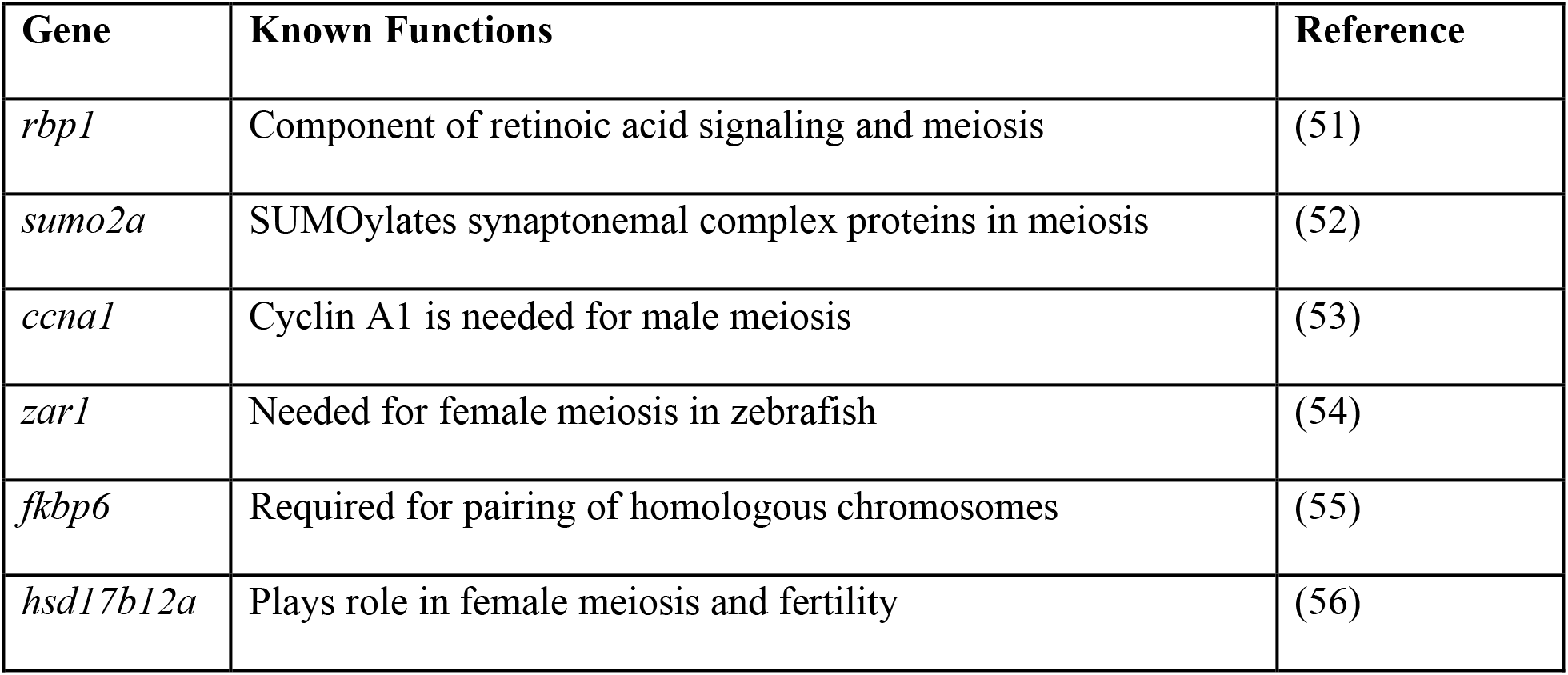
Meiotic genes downregulated in *adad1* mutants.

| Gene | Known Functions | Reference |
| --- | --- | --- |
| *rbp1* | Component of retinoic acid signaling and meiosis | (51) |
| *sumo2a* | SUMOylates synaptonemal complex proteins in meiosis | (52) |
| *ccna1* | Cyclin A1 is needed for male meiosis | (53) |
| *zar1* | Needed for female meiosis in zebrafish | (54) |
| *fkbp6* | Required for pairing of homologous chromosomes | (55) |
| *hsd17b12a* | Plays role in female meiosis and fertility | (56) |

**Table 5.** RNA binding and modification genes downregulated in *adad1* mutants.

| Gene | Known Functions | Reference |
| --- | --- | --- |
| *fkbp6* | Associated with piRNA biogenesis and transposon silencing | (57) |
| *cpeb1b* | Regulates mRNA translation during oocyte maturation | (58) |
| *tdrd6* | Regulates miRNA expression | (59) |
| *rbm11* | Involved in alternate splicing of mRNA | (60) |
| *tsn* | Binds meiotic RNA | (61) |
| *eif4e1b* | Regulates maternal mRNA translation | (62) |
| *magoh* | A component of exon-exon junction complex | (63) |
| *tdrd5* | Binds piRNA and is required for transposon silencing | (64) |

**Fig 10.**
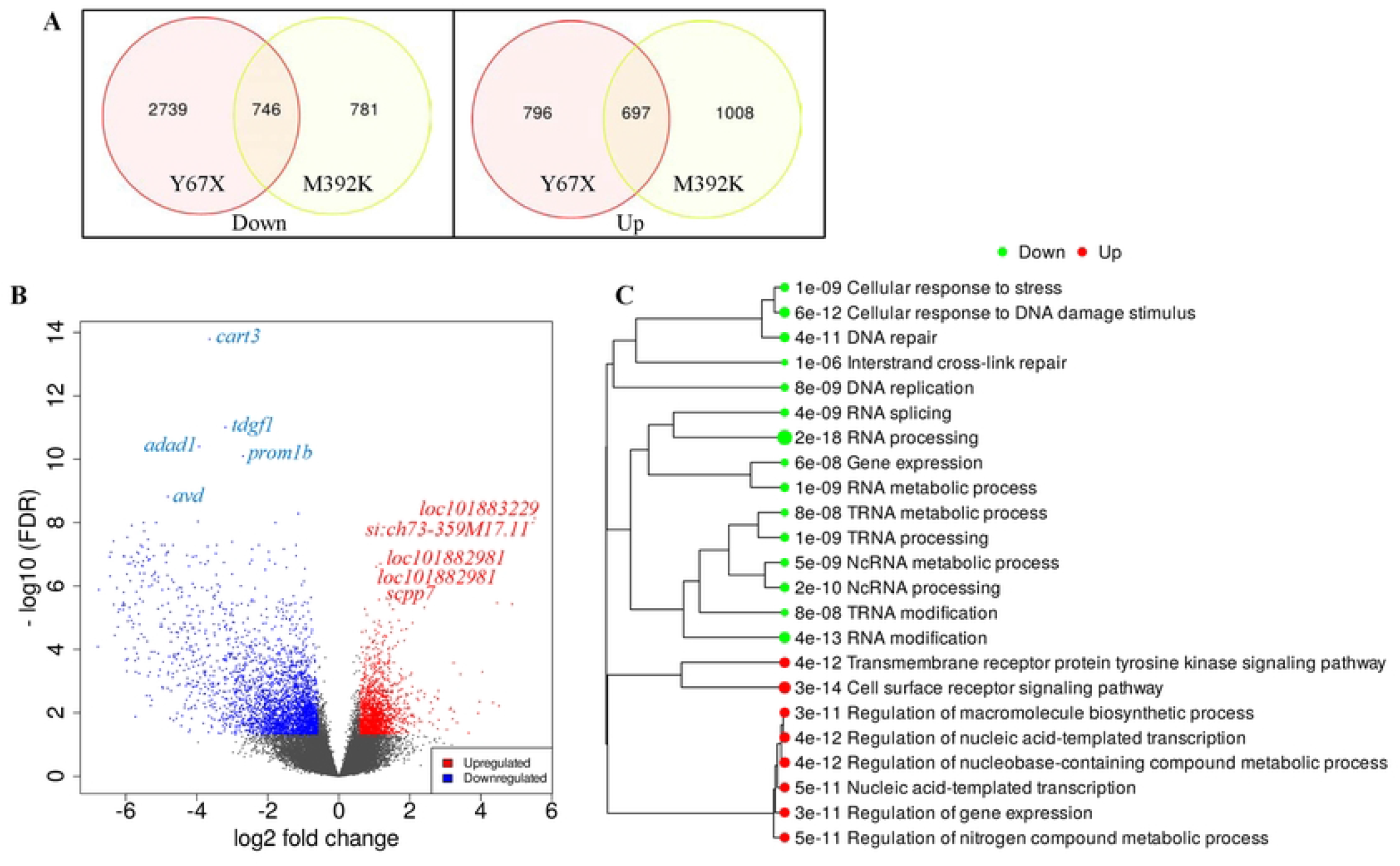
Bulk RNAseq from wild-type and mutant testes. A: Venn diagrams showing numbers of differentially expressed genes in each mutant line using fold change >1.5 and false discovery rate (FDR) <0.05. B: Volcano plot showing genes that were dysregulated in both *adad1* mutant alleles. The top differentially expressed genes defined by lowest adjusted p-values are labeled. C: Gene ontogeny pathway analysis performed on genes that were dysregulated in both *adad1* alleles showing down- and up-regulated pathways in the mutant testes.

## Discussion

In this study, we have shown that the zebrafish Adad1 RNA binding protein has multiple roles in germ cell development. In the testis, Adad1 functions in germline maintenance through regulating SSC function and is also necessary for completion of meiotic prophase I. We propose that Adad1 carries out these roles through regulation of specific RNAs that function in these processes. Our work revealed essential functions of Adad1 in fish, thus increased our understanding of this RNA-binding protein in vertebrates.

### Regulation of spermatogonial stem cells (SSC) by Adad1

We identified *adad1* mutant zebrafish and found that *adad1* functions to maintain the SSCs. We applied an established label retaining cell (LRC) assay to identify quiescence stem cells in zebrafish testes (18). Our LRC experiment showed that Adad1 mutants either lack or have reduced numbers of SSCs compared to their wild-type siblings. Because some mutant testes contained SSCs at the time fish were assayed, we believe that SSCs were initially established in mutants but failed to be maintained. It is also possible that *adad1* mutant SSCs divide less frequently than wild-type SSCs leading to fewer BrdU labeled SSCs after the BrdU treatment. Regardless, SSC function is aberrant in *adad1* mutants. Although *adad1* is expressed in all spermatogonia, proliferation was normal in *adad1* mutant spermatogonia indicating that failure to maintain the SSCs is likely the primary cause of germ cell loss in these mutants.

The establishment and maintenance of SSCs has not been well characterized in fish. *Nanos2*, which is a conserved germline stem cell marker in vertebrates, is expressed in A-type single spermatogonia (A_single_) and spermatogonia in cysts of up to four cells (A_4_) in zebrafish (6–9,65). The *nanos2* gene is required for maintenance and regeneration of germ cells in zebrafish pointing to an essential role in stem cell function (66). Therefore, A_single_ – A_4_ spermatogonia can be presumed to be the stem cells in zebrafish testes. Although Adad1 protein was localized only in the cytoplasm in spermatogonia and spermatocytes, it was also localized in nuclei in A_single_ – A_4_ spermatogonia (Fig 3). These expression data further support a role of Adad1 in regulating zebrafish SSCs and suggests that nuclear Adad1 may be important for stem cell function. Our transcriptomic data points to dysregulation of genes associated with stem cells and pluripotency (Table 2). For instance, *sall4*, which is a well-known SSC marker in mammals, was downregulated in the *adad1* mutant testes (Table 2) (34–36). We also found downregulation of *plppr3b* in mutants, which is a recently identified SSC marker in humans (Table 2) (24,25). Furthermore, the pluripotency factor *nanog*, which is expressed in zebrafish spermatogonia, showed reduced expression in *adad1* mutants (Table 2) (37). Although, *nanog* is generally not associated with spermatogonial cells in mice, it is expressed in spermtogonia of several other fish species, including medaka (*Oryzias latipes*), farmed carp (*Labeo rohita*), and Japanese flounder (*Paralichthys olivaceus*) (67–69). Therefore, it is possible that *nanog* plays a conserved role in regulating spermatogonial cells in fishes. These findings suggest that in the absence of functional Adad1, expression of genes with potential roles in stem cell regulation were disrupted in zebrafish testes, which contributes to germline loss in this vertebrate model.

### Meiosis defects and failure of female development in zebrafish

Because *adad1* mutants initially contained germ cells in larval-juvenile stages, we were able to discover an essential role of *adad1* in meiosis. The *adad1* mutant spermatocytes initiated pairing and synapsis of homologous chromosomes, but failed to complete the process and reach the pachytene stage. We propose that Adad1 binds with RNAs important for meiotic prophase I, thus regulates the process directly, however other possibilities exist, such as regulation of RNAs that indirectly impacts meiosis.

The necessity of Adad1 in meiotic prophase I is further supported by the lack of female development in the mutant lines. Domesticated zebrafish do not have sex chromosomes, and their gonads remain as bipotential until 2-3 wpf (70,71). The bipotential gonads produce meiotic cells that initiate oogenesis. As testes undergo sex differentiation these early oocytes are depleted and spermatogenesis begins (17). Disruption of meiosis in zebrafish gonads leads to male sexual development. This outcome is likely due to failed development of the ovarian follicle in meiotic mutants. In ovaries, the transition from the cyst/nest stage to formation of ovarian follicles occurs around the pachytene stage (72). Thus, defects that disrupt meiotic prophase I typically lead to failed folliculogenesis and male development. Several studies reported a link between meiotic prophase I failure in early oocytes and lack of female development in zebrafish (54,73–79). Zebrafish mutants disrupting genes encoding synaptonemal complex proteins *sycp1*, *sycp2*, and *sycp3* as well as the meiotic cohesion component *smc1b*, developed as 100% sterile males due to failures in meiotic prophase I (73–75,78). Mutations that specifically effect female meiosis, such as mutations in *zar1* and *rad21l1*, failed to develop as female or were male biased, respectively (54,76). In these mutants, males were normal indicating a requirement in female meiosis but not spermatogenesis (54,76). We demonstrated that *adad1* mutant spermatocytes failed to progress beyond the zygotene stage and that ovarian follicles were never formed in the bipotential gonad stage. Therefore, it is likely that *adad1* mutant zebrafish failed to develop as females due to failure in meiotic prophase I.

### RNA-binding proteins and germline maintenance in zebrafish

Post-transcriptional regulation by RNA binding proteins (RBPs) is critical for multiple aspects of germ cell development. (11,12). The functions of RBPs varies and can include translational control, localization, stability, and modifications of RNAs (11,12). Several studies have shown that the conserved germline RBPs Dazl (deleted in azoospermia like), Vasa/Ddx4 (DEAD-box RNA helicase protein 4), Nanos2, and Nanos3 are necessary for germline stem cell establishment or maintenance in zebrafish (8,66,80–82). In *dazl* mutants, the germ cells failed to undergo cystic proliferation and the transition from PGCs to germline stem cells was disrupted leading to a failure to establish germline stem cells (80). Zebrafish *vasa*, *nanos2*, and *nanos3* mutants all established germline stem cells but failed to maintain them (8,66,81,82). The *nanos3* gene is only required for germline stem cell maintenance in ovaries as male mutants were fertile with histologically normal testes (8). By contrast, *dazl*, *vasa*, and *nanos2* are required in both sexes (66,80,81). Here we demonstrated that the zebrafish RBP Adad1 has a critical role in SSC maintenance and progression of meiosis I. We assayed for germline stem cells in juvenile *adad1* mutants using a label retaining cell assay and found that germline stem cells were either reduced or absent by 60 dpf. Thus, similar to *vasa* and *nanos2, adad1* functions in stem cell maintenance in the testis. We could not assess the role of *adad1* in ovarian germline stem cells as oogenesis was never initiated in *adad1* mutants due to the meiotic defects. However, because *adad1* is expressed in both sexes, it may regulate both spermatogonial and oogonial stem cells. Together, these studies indicate that Adad1 is among the multiple RBPs, which are essential for germline stem cell function in zebrafish pointing to critical roles of post-transcriptional processes in germline stem cell regulation.

### Similarities and differences in Adad1 expression and function in vertebrates

The vertebrate specific Adad1 protein has an N-terminal double stranded RNA binding motif (dsRBM) suggesting that it functions in RNA regulation. Adad1 is a member of the Adar (adenosine deaminase acting on RNA) protein family due to the presence of a deaminase domain at its C-terminus (Fig. 1). However, the Adad1 deaminase domain is probably inactive because the amino acid residues known to be required for deaminase activity are not conserved in either zebrafish or mammalian Adad1 (83). Moreover, RNA editing in mouse Adad1 mutant testes was unaffected, indicating that the deaminase domain is not involved in RNA editing (14). The mouse Adad1 protein (also known as Tenr), was first identified due to *in vitro* binding to the *Protamine 1* (*Prm1*) mRNA, however additional target RNAs have not been reported (84). As zebrafish do not have protamines, this RNA target is not conserved across these species (85). Hence, although *adad1* is a predicted RNA regulator, the molecular function of Adad1 remains elusive.

Adad1 is expressed in zebrafish and mammalian germ cells, however differences exist in the precise cell-types in which it is expressed. We found that zebrafish *adad1* is expressed in both testis and ovary, however it is testis specific in humans (proteinatlas.org) and mice (14,86). We found that zebrafish Adad1 is expressed in both spermatogonia and spermatocytes in the testis (Fig. 3), whereas its expression is limited to meiotic and post-meiotic cells in the mammalian testes (14,86). The expression of Adad1 in earlier developmental stages in the zebrafish testis compared to mammals suggests that zebrafish Adad1 regulates early stages of spermatogenesis, as discussed below. Nevertheless, the germline specific expression of Adad1 is conserved from zebrafish to human.

Although both zebrafish and mouse Adad1 have essential roles in male germ cell development, they function in different spermatogenic stages. We demonstrated that loss of *adad1* function triggered germ cell loss due to failure of SSCs maintenance in zebrafish. In addition, these mutants failed to complete meiotic prophase I and therefor never produced sperm. Our results are in harmony with previous studies on mice which reported that *Adad1/Tenr* deficient male mice are infertile due to abnormal sperm development (14,86). However, *Adad1/Tenr* mutant mice only display abnormal sperm morphology and appear to have normal spermatogonial cell maintenance and meiosis. These abnormal sperm have difficulty binding to oocytes and penetrating the zona pellucida resulting in infertility (86). Whether or not zebrafish *adad1* has roles in spermiogenesis similar to those reported in mice cannot be investigated as the earlier defects preclude the ability to investigate spermiogenesis. The human *ADAD1* gene has also been linked to impaired fertility in men. Reduced testicular or sperm *ADAD1* expression was found in male patients and was particularly associated with non-obstructive azoospermia suggesting a role in spermatogenesis (87,88). Overall, these observations suggest that Adad1 functions to regulate spermatogenesis in humans, mice, and zebrafish, although additional roles exist in zebrafish compared to those described in mice.

## Materials and methods

### Zebrafish Lines

Zebrafish were raised and maintained in a recirculating system under standard conditions. Institutional Animal Care and Use Committee approval was attained for all procedures. Zebrafish lines used in this study were: Tue (wild type), *slc34a1a^umb10^, adad1^t30103^, adad1^sa14397^, Tg[ziwi:eGFP]^uc1^* (20), *Tg[ddx4:eGFP]* (19).

The *adad1^t30103^* mutant line was identified in an N-Ethyl-N-Nitrosourea (ENU) mutagenesis screen for defects affecting adult zebrafish gonads, as described previously (89,90). Briefly, wild-type male zebrafish of Tuebingen (Tue) background were treated with ENU. The males were then crossed with females to make F1 families, which were then in-crossed to generate F2 families. Finally, F2 siblings were crossed to obtain homozygous F3 mutants which were raised to adulthood and screened for anomalous gonad phenotypes. The *adad1^t30103^* allele harbors a missense mutation resulting in a Methionine to Lysine change in the protein at amino acid 392, we therefore refer to this mutation as *adad1^M392K^* for simplicity.

The *adad1^sa14397^* nonsense mutant line was generated by the zebrafish mutation project (91) and provided by the Zebrafish International Resource Center (ID: sa14397). This mutant carries a premature stop codon after codon 66, we therefore refer to this mutation as *adad1^Y67X^*.

The *slc34a1a^umb10^* mutants were generated using CRISPR-Cas9 genome editing in the wild-type Tue background. We designed a crRNA (5’-TGTGATCCTGTCCAACCCGG-3’) targeting exon5 of *slc34a1a* using the online tool CRISPRscan (https://www.crisprscan.org/). The crRNA and universal tracrRNA were purchased from Integrated DNA Technologies (IDT). To make the gRNA complex, the crRNA and tracrRNA were mixed and annealed according to the manufacturer’s instructions. The gRNA (5 pg/embryo) and mRNA encoding Cas9 protein (150 pg/embryo, System Biosciences) were injected into one-cell stage embryos. Injected embryos were raised to adulthood and crossed with Tue fish to identify germline mutations. We found F1 embryos carrying multiple different deletions in the gRNA target region from three different parents. However, for further analysis we used the mutant with a 17 bp deletion which caused a premature stop codon after amino acid 195.

### Positional cloning and whole exome sequencing

To identify the heritable mutation of the *t30103* line, we employed whole exome sequencing (WES) as previously described (89) and mapping according to methods described in Bowen et al. (92). Linkage of genomic regions with high mapping scores were tested using SSLP markers and a region on the left arm of chromosome 14 was confirmed to be linked to the gonad phenotype. Two missense mutations were identified within the linked interval, one affecting the *scl34a1a* gene and one affecting the *adad1* gene. Genetic complementation tests between *t30103* and either *slc34a1a^umb10^* or *adad1^sa14397^* revealed that the *adad1* mutation caused the gonad phenotypes. For complementation tests, *adad1* trans-heterozygous fish were generated by crossing *t30103*+/- males with *adad1^sa14397^*+/- females. Similarly, *t30103*+/- males were crossed with *slc34a1a^umb10^*+/- females to establish *t30103/slc34a1a* trans-heterozygous fish. When the fish reached adulthood, fertility tests and gonad histology were performed to test the phenotype.

### Genotyping

To genotype the *adad1^M392K^*, the target region was amplified by primers KS552 and KS553, followed by MslI digestion. The digested PCR product was run on 7% acrylamide gel to distinguish mutant (564 bp) and wild-type (473 bp) bands. Genotyping of the *adad1^Y67X^* mutation was done by PCR amplification using primers KNI203 and KNI204, followed by MseI digestion and electrophoresis on 7% acrylamide gel. The mutant band was 140 bp, whereas the wild type was 166 bp. Primers KS554 and KS555 were used to amplify the region of *slc34a1a* harboring the *slc34a1a^I423N^* variant. This amplicon includes an intronic size polymorphism resulting in 482 bp size mutant and a 582 bp wild-type PCR product. The *slc34a1a* PCR product was run on a 2% agarose gel to separate the mutant and wild-type bands. The *slc34a1a^umb10^* mutation was genotyped by amplifying the gRNA target region using primers KNI201 and KNI202, followed by 7% acrylamide gel electrophoresis to distinguish the mutant band (186 bp) from the wild type (203 bp). Sanger sequencing was done on PCR products to confirm all the mutations. The name and sequence of the genotyping primers is listed in S1 Table.

### Fertility tests

The fertility test of *adad1^M392K/Y67X^* trans-heterozygotes (N=6) and *adad1^Y67X^* homozygous mutants (N=3) were done by pairing mutant males with wild-type Tue females, since the mutants were all male phenotypically. The following morning, embryos were collected and monitored under the dissecting microscope to see the developmental stage. Embryos were considered to be fertilized if they had undergone normal appearing cell divisions. Eggs that did not appear to be developing were raised up to 24 hours to confirm that they were not fertilized, by which time all of them died and became cloudy.

### Histology

Histology was performed as previously described (93). Briefly, fish were euthanized by tricaine overdose and the torso was isolated and fixed in Bouin’s solution overnight at room temperature. Following several dehydration steps, the fixed tissues were embedded in paraffin for sectioning. A rotating microtome was used to obtain 5 μm thick sections and the sections were stained following modified Harris’s hematoxylin-eosin staining protocol. After staining, the tissue was covered using Permount as a mounting medium.

### RT-PCR

Total RNA was isolated from zebrafish testes or ovaries using TRIzol-reagent, following the manufacture’s protocol (Fisher Scientific). Isolated RNA was treated with TURBO DNase (Fisher Scientific) to remove any genomic DNA contamination and cleaned via LiCl precipitation. The first strand cDNA was synthesized using oligo-dT primers and AMV-Reverse Transcriptase (New England Biolabs). The *ziwi* gene is expressed in the germ cells of both sexes and was used as germline control. *Rpl13a* expression was used as a control for other tissues. The primers used for RT-PCR are listed in S3 Table. To generate a zebrafish testes devoid of germ cells for RT-PCR using male somatic gonadal tissue, we injected a morpholino targeting the *dead end* gene into 1-cell embryos, as previously described (94).

### *In situ* hybridization (ISH)

To make the *adad1* ISH probe, a 341 bp cDNA fragment was amplified using *adad1* RT-PCR primers KNI207 and KNI208. The cDNA was sub-cloned into pGEMT easy vector (Promega), and Sanger sequencing was done to confirm the right insert. The sense and anti-sense probes were generated using a DIG RNA labelling kit (Roche). ISH on 4% paraformaldehyde (PFA) fixed, paraffin embedded tissue sections was performed as previously described (89) To visualize the probe, alkaline phosphatase conjugated anti-DIG antibody (Roche), and BM purple substrate were used.

### Adad1 antibody generation and western blotting

Polyclonal antibodies were raised in rabbits to the peptide sequence KSYLERKGYGQWVEKPPISDHFSI by Pacific Biosciences, CA. Antibodies used in this study were purified by affinity chromatography. To test the antibody through western blot, we isolated total protein from 45 dpf *adad1^Y67X^* mutant and wild-type testes in RIPA buffer (Thermo Fisher Scientific). After denaturation through heat and 100 mM DTT, the samples were run on 10% SDS-PAGE and transferred onto a PVDF membrane. The membrane was blocked in 5% milk in TBST for 1 hour at room temperature. Primary antibodies were diluted in TBST as follows: rabbit Adad1 at 1:500, rabbit Vasa at 1:2000 (95), mouse α-Tubulin at 1:1000 (T9026, Sigma-Aldrich). The membrane was incubated in primary antibodies overnight at 4°C. After 4 x 5 minutes washes in TBST, incubation in the secondary antibodies (Li-COR Biosciences) were done for 1 hour at room temperature. Finally, the membrane was washed 4 x 5 minutes in TBST before imaging in the Odyssey CLx instrument (Li-COR Biosciences).

### BrdU incorporation experiments

For cell proliferation assays, 45 dpf mutant and wild-type fish were kept in BrdU solution (3 mg/mL) overnight at 27°C. The next morning, the BrdU solution was removed, and fish were kept in fresh water for 6 hours before euthanizing. The torsos were fixed in 4% PFA, embedded and sectioned in paraffin, and processed for immunofluorescence (IF). For label retaining cell (LRC) assays, 32 dpf non-genotyped fish were kept in BrdU solution (3 mg/mL) for three consecutive nights. Between treatments the fish were kept in fresh fish water to avoid BrdU toxicity. After a 25 days chase period, tissue was collected for genotyping and the mutant and wild-type fish were fixed and processed for IF as described below.

### Immunofluorescence and cell death assay

The process of tissue fixation and antigen retrieval for IF was done as previously described (74). TUNEL staining on 60 dpf paraffin embedded testes sections were performed using a fluorescein *In Situ* Cell Death Detection Kit (11684795910, Roche). Immunolabelling with Vasa antibody was done before the TUNEL staining. Primary antibodies were diluted in the blocking buffer (1% goat serum in PBST) as follows: rabbit Adad1 at 1:500, chicken Vasa at 1:2000 (79), rabbit Sycp3 at 1:200 (NB300-232SS, Novus Biologicals), mouse BrdU at 1:200 (5292S, Cell Signaling Technology), rabbit Caspase-3 at 1:1000 (C8487, Sigma-Aldrich). The testes sections were incubated overnight with the primary antibodies at 4°C. Incubation with appropriate secondary antibody (Thermo Fisher Scientific) was done at room temperature for 1 hour. Following 10 minutes incubation with nuclear stain DAPI, sections were washed 3×5 minutes in PBST and 1×5 minutes in miliQ water. Finally, the sections were covered with Fluoroshield (Sigma-Aldrich) and kept at 4°C before imaging.

### Chromosome spreads and staining

Meiotic chromosome spreads from 8 wpf mutant and wild-type testes were prepared as previously described (74,96). Telomere staining was done using a TelC-Cy3 probe (PNA Bio) following a previously described method (97). Antibody labeling of chromosome spreads was done as described above.

### Imaging

We used a confocal laser scanning microscope (Zeiss LSM 880) to capture images of the IF and telomere staining. The images were further analyzed through Fiji/ImageJ. To quantify cell numbers, we used 3 sections for each biological replicate (6-8 replicates for each genotype). Student’s t-test (p<0.05) was employed to see whether difference between the mutant and wild-type samples were significant or not.

### RNA sequencing and data analysis

Total RNA was isolated from 45 dpf testes using the RNeasy micro kit (Qiagen). Since the testes were very small at this age, 5 testes were pooled together for each biological replicate (4 biological replicates for each genotype). Isolated RNA was sent to Genewiz at Azenta Life Sciences, NJ for library preparation with poly-A selection method and 150-bp paired end sequencing on an Illumina HiSeq platform. We assessed the quality of the raw sequence data using FastQC, low quality and adapter sequences were trimmed with TrimGalore. Trimmed reads were aligned to the GRCz11 assembly of the zebrafish genome (Ensembl) using STAR and the gene annotations published by the Lawson lab (98). Reads were counted using HTseq and analyzed for differential gene expression (DEG) and gene ontology (GO) using the iDEP v95 pipeline (99), with a DeSeq2 cut off value for fold change of >1.5, and false discovery rate (FDR) of less than 0.05.

## Acknowledgements

We would like to thank Sean Burgess (UC Davis) for providing Sycp1 and Vasa (raised in chicken) antibodies. We are also grateful to Holger Knaut (NYU) for generously giving Vasa antibody (raised in rabbit). Special thanks to Leo Harris for the zebrafish drawings used in figures 8 and 9. The *adad1^sa14397^* mutant line was provided by the International Zebrafish Resource Center (ZIRC). The mutagenesis screen in which the *adad1^t30103^* mutant was isolated was supported by the European Commissions, ZF-MODELS 6th framework program (EC Contract LSHG-CT-2003-503496).

---

**Fig S1. Genomic DNA sequence from *adad1* nonsense and missense alleles.** The nonsense mutation (*sa14397*) is located in exon 3 where a T<A mutation caused a premature stop codon (left panel). The missense mutation (*t30103*) is in exon 9 where a T<A mutation resulted in a Methionine to Lysine residue change (right panel).

**Fig S2. Testis expression and phenotype of *slc34a1a*.** The *scl34a1a^umb10^* mutation is a 17 bp deletion (top panel). The guide-RNA target sequence, including the PAM, is highlighted in yellow. RT-PCR detected *slc34a1a* expression in adult testes (left panel). Homozygous *slc34a1a^umb10^* mutants exhibited normal testes histology. Scale bar: 50 μm.

**Fig S3. Zebrafish Adad1 antibodies specifically label Adad1 protein.** A-D: Immunolabeling of Adad1 antibody on 60 dpf *adad1^Y67X^* mutant testes. No protein was detected as expected since the antibody epitope is at the C-terminus of the protein (A). Co-labeling with Vasa showed the presence of germ cells in the mutant testis (B). E: Western blot showing that the antibody recognized the 62 kDa size Adad1 protein in wild-type testes extracts but not in testes extracts from *adad1^Y67X^* mutants. The wild-type sample was loaded in two lanes: the left lane had twice the volume as the right lane. Vasa was detected in the mutant sample indicating that germ cells were present. Scale bar: 20 μm.

